# Meta(xylem)morphoses: Root vascular patterning is dynamic and convergent in temperate grasses

**DOI:** 10.64898/2026.11.19.767855

**Authors:** Dylan H Jones, Maria Schön, Darshan Damodaran, Corinna Trautewig, Juan Baca Cabrera, Alexander Ware, Michael Melzer, Matias Schierenbeck, Dongxue Zhao, Raul Gimenez, Guilliaume Lobet, Bipin Pandey, Valentin Couvreur, Malcolm Bennett, Daniel Rodriguez, Hannah M Schneider

## Abstract

In cereal roots the metaxylem vessels are the main channels of axial water transport. Metaxylem size and number are considered constant traits characterising different root classes. Here we show that metaxylem number is progressively remodelled along root axes, producing mature roots with vascular anatomy differing by position. At initiation, adventitious wheat root tips produce multiple files of metaxylem, and then progressively fewer, until a single central metaxylem file exists, resembling the vascular anatomy of the seminal roots. This pattern was observed in wheat, barley, rye, oat, and *Brachypodium*, but not in maize or sorghum. We investigated the developmental regulation of this phenomenon by creating a cell-resolved spatial transcriptomic atlas of wheat adventitious root tips in these different metaxylem production states and integrated this with bulk RNA-seq analysis of whole root tips. We developed a pipeline integrating these complementary transcriptomic datasets to reveal differentially expressed genes that included candidates controlling meristem maintenance, vascular patterning, and metaxylem wall lignification. Hydraulic modelling revealed changes in vessel number can alter the distribution of hydraulic pressure levels in the root, while maintaining or increasing the capacity of roots to transport water. We propose that this developmental transition preconfigures different regions of the root for their changing hydraulic environments, providing a mechanism to balance hydraulic robustness and efficiency over the root life cycle.

## Introduction

Water transport is fundamental to plant growth, survival, and productivity. In cereals, the transport efficiency of water and mineral nutrients from the soil to the plant aerial tissue is determined not only by the size and architecture of the root system, but also by the cellular organization of its vascular tissues (Jones et al., 2025). Despite this central role in plant physiology, root vascular systems remain substantially less studied than their aerial counterparts, particularly in mature field-grown plants where recovering intact root systems is technically challenging (Ryan et al. 2016). Consequently, how vascular organisation develops and is maintained across the lifespan of cereal roots remains poorly understood.

The principal conduits for axial water transport in cereal roots are the metaxylem vessels, which are positioned within the stele, the central vasculature cylinder. Once differentiated, metaxylem vessels undergo programmed cell death to leave continuous files of hollow, lignified cells that form vessels responsible for water transport from the roots to the aerial tissue. Lignification enhances the cell files physical strength, allowing them to withstand the negative pressures generated by transpirational pull (Zimmermann et al., 2002). Axial conductance (Kx) is the sum of the conductances of the individual vessels which as this scales to the fourth power of vessel radius [as per the Hagen-Poiuseuille law (Tyree and Zimmermann, 2002)], small perturbations in vessel size can have large consequences for water transport capacity. Therefore, a single wide vessel has a far greater transport capacity (Sanderson et al., 1988; Steudle and Peterson, 1998) than multiple narrower vessels with the same total cross-sectional area. Alternatively narrower vessels can present an advantage under water limitation, slowing water use to delay soil water depletion and reducing the risk of cavitation (Harrison Day et al., 2023). Variation in metaxylem size and number have accordingly been linked to water use, drought tolerance, and yield across various cereals under controlled (Kadam et al., 2015; Klein et al., 2020; Strock et al., 2021; Clément et al., 2022; Affortit et al., 2024) and field conditions.

Unlike dicotyledonous roots which develop additional vascular files through secondary growth, cereal roots lack a vascular cambium and build their metaxylem files entirely through primary growth. Metaxylem files are patterned close to the root apical meristem, after which their number is developmentally stable, while many root morphological features such as lateral root patterning, secondary cell wall modification, and cortical senescence (Van Norman et al., 2013; Jones et al., 2025; McCahill et al., 2025; Dhar et al., 2026) change with tissue age. This fixed post-developmental pattern of the metaxylem can therefore be read along the root axis as a stable diary of the developmental decisions made at the root tip during growth. While many studies rely on anatomical sampling at a single position to infer genetic regulation or physiological impact (Koehler et al., 2025), this static approach misses the broader temporal dynamics of tissue differentiation. Uncovering these dynamic pattern variations over the course of root growth reveals how plants actively reprogram cell fates long after early embryonic patterning is complete. Consequently, the functional consequences of pattern variations occurring over the course of root growth remain largely unconsidered.

Grass root systems are a particularly informative system in which to address this question as they comprise several different classes of axial roots that arise in sequence. Seminal roots emerge from the embryo and support early seedling establishment (Hochholdinger et al., 2018), followed by adventitious roots which emerge in acropetal sequence from stem nodes (Hochholdinger et al., 2004; Liu et al., 2020; Schneider et al., 2021), and eventually make up the majority of the root system in older plants. These classes can differ significantly in anatomy, growth rate, lifespan and function (Liu et al., 2020; Koehler et al., 2025), whilst further differences can be made between adventitious roots originating from different stem nodes. However, these observations of difference are typically made at a single comparable sampling position. Maintaining vascular continuity during growth requires developmental coordination and necessitates conductive elements remain connected despite continuous cell division, differentiation, and elongation occurring at the root apex. Considerable progress has been made in understanding the molecular regulation of vascular patterning and continuity in *Arabidopsis*, where auxin transport, canalisation, and programmed cell death coordinate the formation of connected vascular networks (De Rybel et al., 2016; Berleth et al., 2000; Scarpella, 2017). In comparison, far less is known about how metaxylem continuity is established and maintained in grass roots, and still less in the mature adventitious roots which dominate the later root network.

Here we surveyed root anatomy in intact mature root systems across multiple grass and cereal species, sampling different root classes and positions along the axis. We describe a previously unrecognised developmental transition during which adventitious roots progressively re-pattern their metaxylem during growth. This coordinated reduction in metaxylem files drives an anatomical convergence on a vascular anatomical phenotype common to adventitious and seminal roots. This transition phenomenon was conserved across a range of temperate grasses surveyed (wheat, barley, rye, oat, and brome) but was absent in panicoid grasses (maize and sorghum). To investigate its developmental basis, we utilised cell-resolved spatial transcriptomics to create an atlas of adventitious root tip development at different ages and combined this with bulk RNA sequencing to detect root class and age interactions. To understand the bearing of this vascular repatterning on root physiology we utilised hydraulic modelling to understand functional dynamics at the local and whole organ levels. Together these analyses reveal an unexpected phase of vascular remodelling in adventitious roots of grasses and provide new insight into how vascular continuity is maintained during root development.

## Materials and Methods

### Plant material and growth conditions

Plants were grown in three experimental systems: hydroponic culture, soil-filled mesocosms and field trials. The species, genotypes and system used for each experiment are summarised in Table S1.

### Hydroponic culture

Seeds were surface-disinfected with 10% NaClO for 5 min, rinsed three times in distilled water and germinated on filter paper moistened with tap water in a controlled-environment chamber at 22 °C, 60% relative humidity and a 16 h light/8 h dark cycle with a photosynthetic photon flux density of 400 µmol photons m⁻² s⁻¹ (ISTA, 2018). After seven days, fifteen uniform seedlings per genotype were transplanted into 75.5 L hydroponic tanks (Rubbermaid Brute; 38.3 × 70.7 × 44.1 cm) containing a continuously aerated nutrient solution that was replaced every two weeks (Schierenbeck et al., 2026). Plants were maintained at 17 °C day and 15 °C night under a 16 h photoperiod (900 µmol photons m⁻² s⁻¹). For the bulk RNA-seq and spatial transcriptomic experiments and the developmental time series, plantings were staggered across four dates at two-week intervals, so that young and old adventitious and seminal roots were available at a single harvest.

### Soil mesocosms

Seeds were pre-germinated on paper rolls for five days, after which two seedlings were transplanted per column and later thinned to one after establishment. Plants were grown in 30 L PVC mesocosms (110 cm length, 23 cm internal diameter) lined with a plastic sleeve and perforated at the base for drainage, at 25/16 °C under a 16/8 h photoperiod and a photosynthetic photon flux density of 830 µmol photons m⁻² s⁻¹ for 48 days. The substrate (bulk density 0.86 g cm⁻³) consisted of 50% medium sand (particle size 0.20–0.63 mm), 20% agricultural Chernozem A-horizon sieved soil of sandy clay loam texture (Altermann et al. 2005;USDA, 2017) from the experimental fields of IPK Gatersleben (Germany), 15% perlite (Perligrain Classic, 0.1–6 mm) and 15% vermiculite (Floragard, 1–2 mm), amended with a controlled-release fertiliser (Osmocote 5 3-4M, 16N–8P–12K–2.2% MgO plus micronutrients) at 5 g L⁻¹. Nutrient availability was further supplemented with nutrient solution as described by Schierenbeck et al. (2026): maize received 825 mL per plant and temperate cereals 525 mL per plant, each split into five applications. Plants were watered twice weekly according to crop demand to maintain non-limiting water conditions, giving a cumulative irrigation volume of 4.17 L per plant for maize and 3.67 L per plant for temperate cereals. Before harvest, irrigation was withheld for approximately four days to allow drainage; columns were then dismantled, root systems washed free of substrate, and stored in 50% ethanol until imaging.

### Field trials

The sorghum trial was conducted at Warwick, Queensland, Australia (28°12′46″S, 152°06′14″E) during the 2024/25 summer season, comprising 250 plots (5 × 3 m) of 53 hybrids at 0.75 m row spacing and two plant population densities (6 and 12 plants m⁻²), on a deep black cracking clay (Vertosol; Isbell, 2021), sown on 20 December 2024. The wheat trial was conducted at Gatton Research Station, Queensland (27°32′44″S, 152°19′54″E) during the 2024 winter season, as a factorial combination of two plant populations (60 and 120 plants m⁻²) and 36 cultivars (13 commercial varieties and 21 breeding lines) in a split-plot design with four replicate blocks, on black and grey cracking clays (Isbell, 2021; Powell, 1982), sown on 15 May 2024 into soil near field capacity. Root sampling for both trials is described below.

### Root sampling and anatomical phenotyping

Root systems from mesocosm-grown plants were separated into classes by origin; seminal and nodal adventitious roots were distinguished in all species, and in maize and sorghum the embryonic primary root was distinguished from the seminal roots. Each root was divided into 10 cm segments (5 cm for wheat), and a 2 cm subsegment was taken from the midpoint of each segment. These sections were imaged, annotated and measured with the Rapid Anatomics Tool (RAT; Jones et al., 2025) following region-of-interest definition in FĲI (ImageJ 1.54; Schindelin et al., 2012), yielding metaxylem file number, individual and total metaxylem area, stele area and complexity (the metaxylem perimeter-to-area ratio normalised by that of the stele); these measurements underlie Figure 1D and Figure 2. Field root systems were sampled by soil coring (sorghum, 4.4 cm auger to 1 m, two on-row cores per plot; wheat, 5 cm auger to 0.9 m, one core adjacent to a representative plant per plot); cores were cut into 10 cm depths, washed and, for wheat, a 2 cm subsegment of the largest-diameter root taken from alternate depths (0–10, 20–30, 40–50, 60–70, 80–90 cm). The three thickest roots per depth were retained in 60–70% ethanol and pooled by depth (wheat, n = 12 cores; sorghum, n = 6 cores) before imaging as above using the RAT.

**Figure 1.**
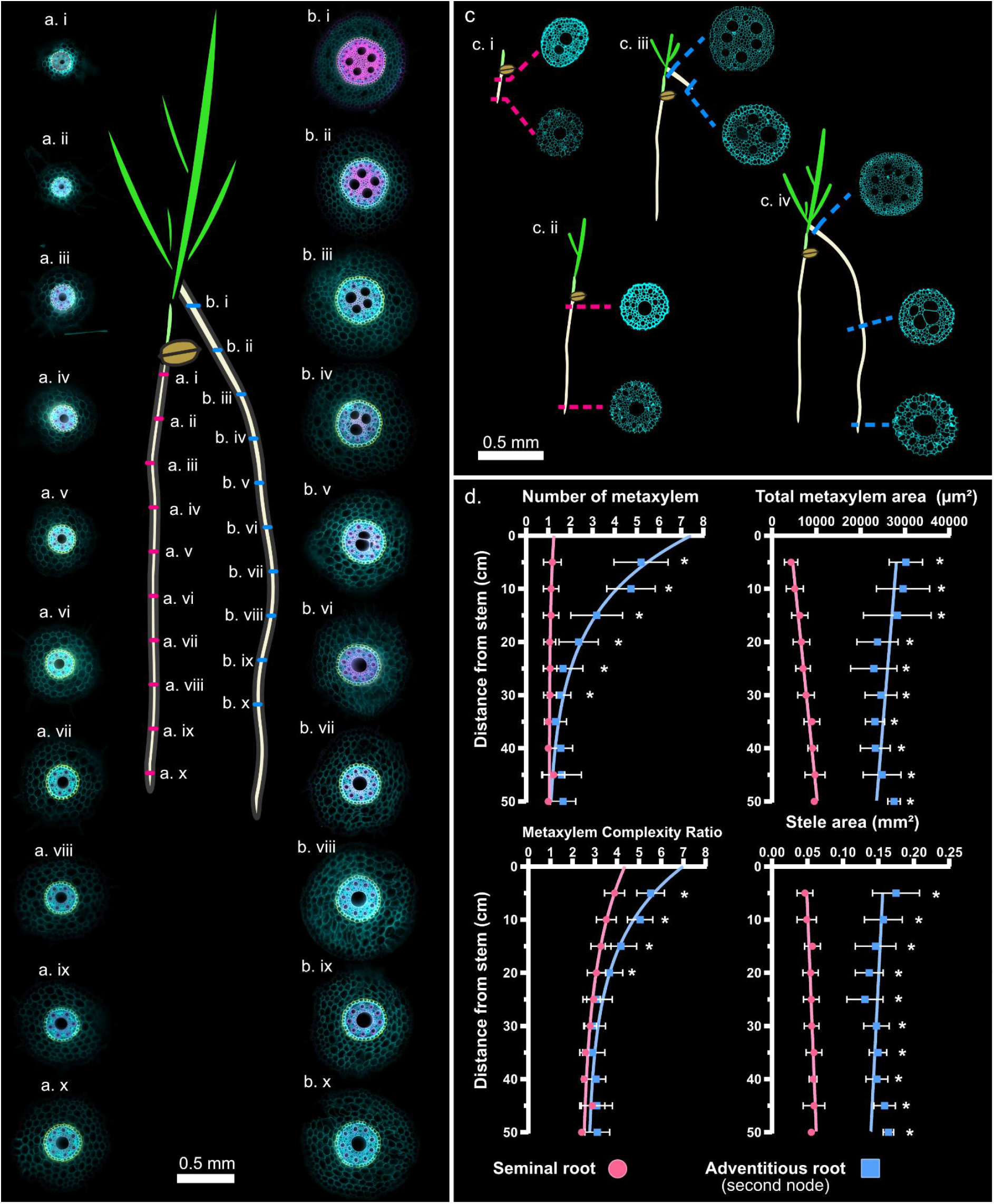
Metaxylem anatomy converges between adventitious and seminal roots in wheat. (a, b) Root class origins and sampling positions, with the transverse section imaged at each. Roots were divided into 5 cm segments and one section taken from each midpoint (± 1 cm); sections i–x span ∼2.5 to ∼47.5 cm from the stem in (a) seminal (pink) and (b) second-node adventitious (blue) roots. Stained for callose/cellulose (cyan), lignin (magenta) and suberin (yellow). [Scale bar, 0.5 mm] (c) Stele cropped sections in roots of different lengths, each sampled within 1 cm of the base and 1 mm of the tip: (c.i) 8 cm and (c.ii) 30 cm seminal roots; (c.iii) 8 cm and (c.iv) 35 cm adventitious roots, the latter also sectioned 17–18 cm from the base. (D) Metaxylem file number, total metaxylem area, complexity and stele area versus distance from the stem, for adventitious (blue) and seminal (pink) roots; complexity ratio is the metaxylem perimeter-to-area ratio normalised by that of the stele. Asterisks, significant difference between classes at a position (two-way ANOVA, Tukey’s multiple comparison correction). Error bars, SD; n = 16.

**Figure 2.**
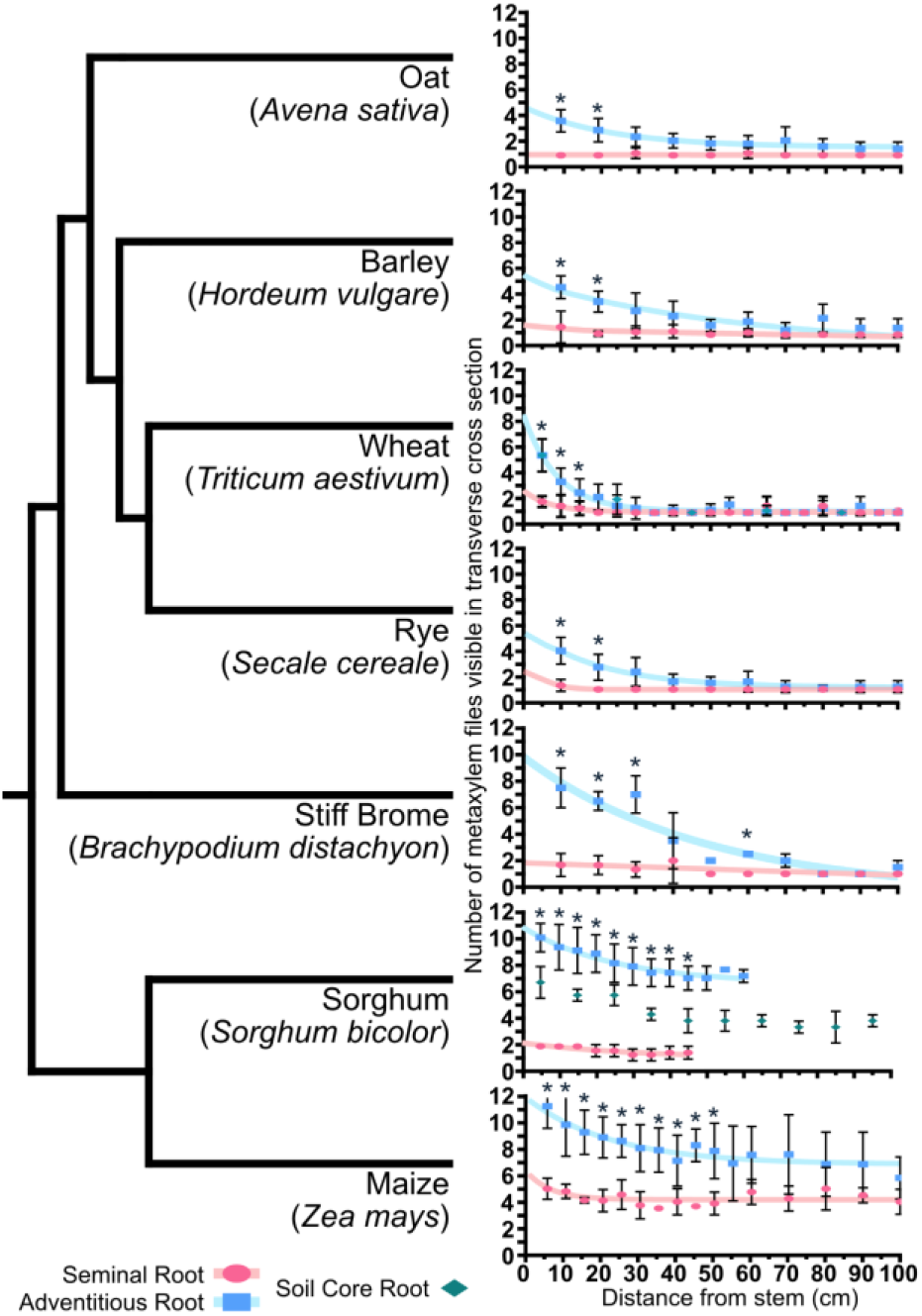
Adventitious root metaxylem file number converges on that of seminal roots with distance from the base across multiple grasses. Cladogram of the seven grasses surveyed, each paired with metaxylem file number versus distance from the stem for seminal (pink), second-node adventitious (blue) and, where sampled, soil-core (green) roots. Mesocosm-grown plants, two pooled genotypes per species, n = 6; field plants sampled from soil cores, the three thickest roots per core from two cores per plot across four plots and three genotypes, pooled by depth (wheat n = 12, sorghum n = 4). Asterisks, significant difference between adventitious and seminal roots at a position (two-way ANOVA, Tukey correction). Cladogram adapted from Grass Phylogeny Working Group III (2024)

### Histology and light microscopy

For the representative sections in Figure 1a and b, fine hand sections of seminal and second-whorl nodal roots (5 cm segments, midpoint ± 1 cm) were fixed in 70% ethanol, 10% acetic acid and 20% water. All staining was performed in the dark with gentle shaking. To visualise suberin and lignin, sections were stained with Basic Fuchsin (Thermo Scientific; 0.2% w/v in 70% ethanol) for 10 min and washed twice in 100% ethanol, then with Fluorol Yellow 088 (ChemCruz; 0.01% w/v in methanol) for 5 min and washed in 50% ethanol, and counterstained with Calcofluor White (Sigma-Aldrich) for 30 min before rinsing thoroughly in MilliQ water. Sections were imaged the same day on a Keyence BZ-X1000 with a 10× objective, acquiring Basic Fuchsin in the TRITC channel (excitation 545 nm, emission 570–640 nm), Fluorol Yellow in the GFP channel (470 nm, 500–550 nm) and Calcofluor White in the DAPI channel (360 nm, 435–485 nm). For the high-resolution sections in Figure 1C, four-to-five-millimetre segments from seminal and adventitious roots were fixed by microwave-assisted aldehyde and osmium tetroxide fixation (2% glutaraldehyde and 2% paraformaldehyde in 0.05 M cacodylate buffer, pH 7.3, followed by 1% osmium tetroxide), dehydrated through a graded acetone series and embedded in Spurr’s resin using a PELCO BioWave 34700 microwave processor (Ted Pella, Redding, CA, USA; full schedule in Table S2); sectioning and light microscopy followed Kavka et al. (2024), images were saved in 8-bit, inverted, and a Cyan LUT was applied and images were cropped to the stele.

### Spatial transcriptomics

*sample preparation.* Young (AY; 5–10 cm, 32 days after germination) and old (AO; 30–40 cm, 46 days) node-2 and node-3 adventitious root tips were collected from hydroponically grown wheat, trimmed to 3 cm and then to 8 mm, rinsed and embedded in optimal cutting temperature compound (OCT). To maximise the number of tips per section, 16 (AY) and 12 (AO) root tips were arrayed in a custom three-dimensionally printed insert (0.2 mm walls) within a 7 mm Tissue-Tek cryomould in 1 mm OCT layers of three to four tips; layers were frozen on dry ice and assembled into a single compound block.

### Capture, library preparation and sequencing

Stereo-seq capture, library preparation and sequencing were performed as a commercial service by BGI Genomics (Shenzhen, China) following the standard STOmics Stereo-seq protocol, as was the bulk RNA-seq library preparation and sequencing (below). Compound blocks were sectioned transversely from the face furthest from the tips in 10 µm increments, selecting a section immediately proximal to the meristematic region of the root tip, where the root cap was visible and metaxylem were present but not yet fully differentiated and still nucleated, and captured on Stereo-seq chips (500 nm bead-to-bead distance).

### Spatial data processing

Reads were processed with the SAW pipeline (Gong et al., 2024): coordinate and molecular identifiers were taken from read 1 (nucleotides 1–25 and 26–35 respectively) and the cDNA from read 2, coordinate identifiers were mapped to the chip’s spatial coordinates allowing one mismatch, and reads with an N or low-quality base in the molecular identifier or a poly-A body were discarded. Retained reads were aligned to the IWGSC RefSeq v2.1 wheat genome (Zhu et al., 2021) with the STAR-based aligner within SAW (Dobin et al., 2013); uniquely mapped reads were annotated to genes, molecular identifiers sharing a coordinate and gene were collapsed, and a spatial gene-expression matrix was generated for each chip.

*Cell segmentation and annotation.* Four sections were selected per chip. For each, a maximum-projection image of the nuclear (DAPI) and cell-wall (Fluorescent Brightener 28) stains was co-registered with the Stereo-seq expression matrix, and individual cells were annotated manually in FĲI by drawing region-of-interest polygons on the cell-wall image; polygons were converted to coordinate arrays and captured transcripts assigned to the enclosing cell. Cells were assigned to one of 15 tissue classes, yielding 6,685 annotated cells across eight sections.

### Clustering and marker analysis

In Scanpy (v1.10.3; Wolf et al., 2018), per-cell counts were normalised to counts per 10,000 and log-transformed, 3,000 highly variable genes were selected (seurat_v3) and principal component analysis was performed on the scaled data (50 components). Batch effects between chips and sections were corrected with Harmony (harmonypy v0.0.10; Korsunsky et al., 2019), a neighbourhood graph was constructed (50 neighbours over 50 principal components), cells were clustered with the Leiden algorithm (leidenalg v0.10.2; Traag et al., 2019) at resolution 1.4, and visualised by UMAP (umap-learn v0.5.6; McInnes et al., 2018). Tissue-enriched markers were identified using scikit-learn (Pedregosa et al., 2011) by a one-versus-rest Wilcoxon rank-sum test of each tissue against all others, with the area under the receiver-operating curve derived from the U statistic, retaining genes with AUC ≥ 0.70 and a Benjamini–Hochberg-adjusted p < 0.05. The response of each gene to root-tip age was tested within tissue by a two-sided Mann–Whitney U test between young and old cells and summarised as the log_2_ fold change (young versus old); tissue-relative expression was min–max scaled across the 15 tissues for display.

### Bulk RNA-seq

For each replicate, a pool of ten root tips from different plants was collected, with each of the four sample groups (seminal young, seminal old, adventitious young and adventitious old) represented by four biological replicates. Total RNA was extracted with TRIzol (Invitrogen), genomic DNA removed with the TURBO DNA-free Kit (Invitrogen) and RNA quantified by Qubit (Invitrogen); library preparation and sequencing were performed by BGI Genomics (above). Clean reads were aligned by BGI to the IWGSC RefSeq v1.1 wheat genome (IWGSC, 2018) with HISAT2 (v2.2.1; Kim et al., 2019) and gene-level counts summarised with featureCounts (Rsubread v2.18; Liao et al., 2014); gene identifiers were subsequently converted to RefSeq v2.1 (Zhu et al., 2021), so that the bulk and spatial datasets shared a common annotation. Differential expression was assessed with DESeq2 (v1.46; Love et al., 2014); genes were called differentially expressed at a Benjamini– Hochberg-adjusted p < 0.05 with no fold-change threshold, association with metaxylem file number instead being defined by concordant direction of change across the two target contrasts. The target contrasts were adventitious young versus adventitious old (AY/AO) and adventitious young versus seminal young (AY/SY), with adventitious old versus seminal old (AO/SO) and seminal young versus seminal old (SY/SO) retained as exclusion contrasts.

### Candidate gene filtering

Genes associated with metaxylem convergence were selected by successive filtering (Figure 3a, b). Of the 82,074 genes detected in common, those differentially expressed and concordant in direction across the AY/AO and AY/SY contrasts, and not driven by the AO/SO or SY/SO exclusion contrasts, were retained (5,093 genes). This set was filtered against the spatial data: genes more highly expressed in metaxylem than in a pseudobulk of all other tissues were retained (1,255 genes); of these, genes concordant in direction between the bulk contrast and the young-adventitious metaxylem pseudobulk were retained (739 genes); and finally genes with their highest per-cell mean expression in metaxylem were retained (448 genes). In parallel, the same spatial filtering applied to genes also significantly differentially expressed in the per-section spatial pseudobulk yielded a stringent set of five candidates.

**Figure 3.**
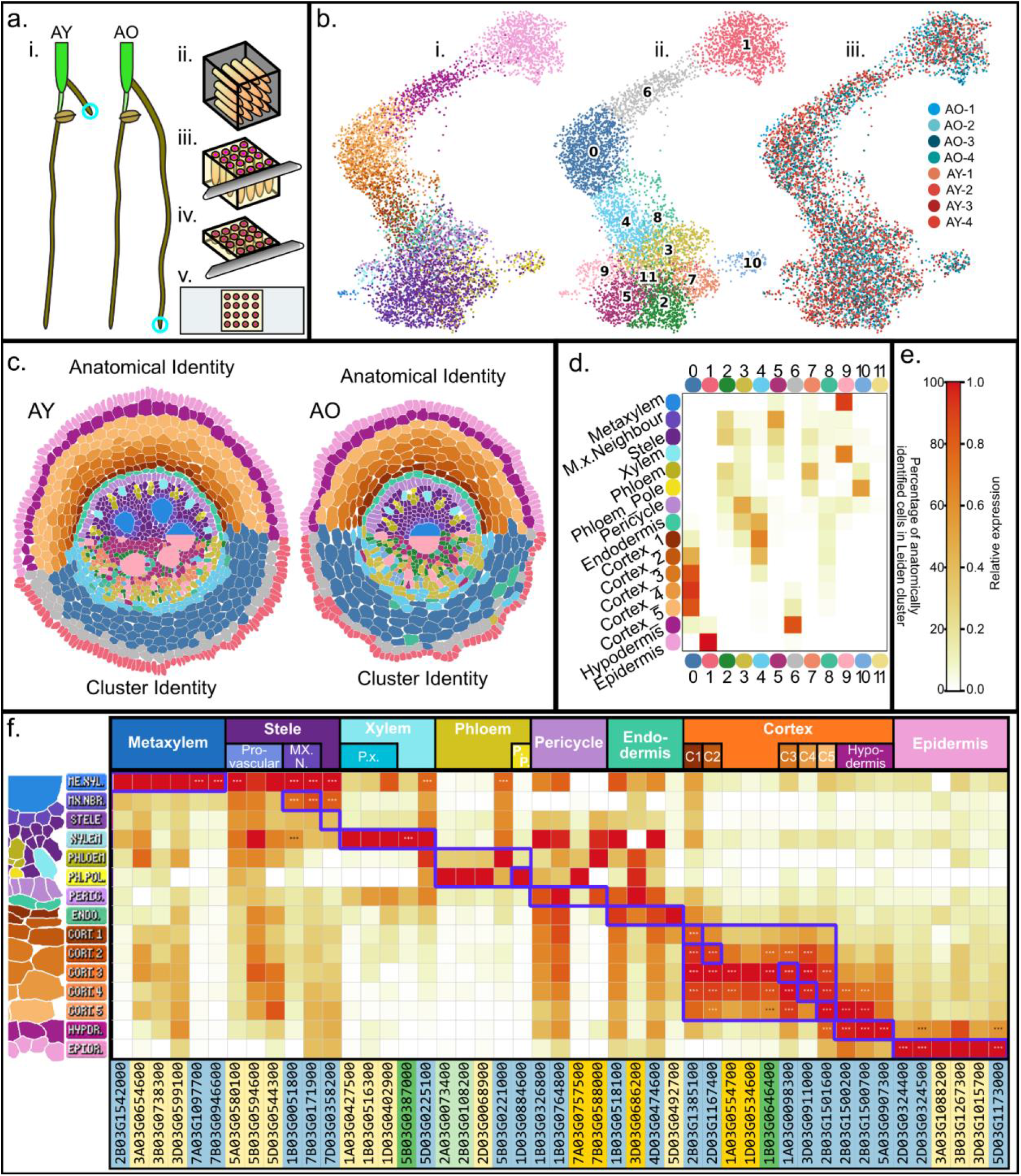
Cell-annotated spatial RNA-seq shows high concordance between manually and statistically defined cell groups expressing known wheat marker genes. (a) Sampling and section preparation: (a.i) Adventitious Young (AY; 32 days after germination, 5–10 cm, n = 16) and Adventitious Old (AO; 46 days, 35–40 cm, n = 12) second-node tips were excised, trimmed to 5 mm, (a.ii) OCT-embedded and (a.iii, a.iv) sectioned transversely from the base to just above the quiescent centre, where metaxylem were present but undifferentiated and still nucleated; (a.v) captured on Stereo-seq chips (one per age). (b) UMAP of 6,685 annotated cells coloured by (b.i) tissue identity, (b.ii) Leiden cluster (resolution 1.4) and (b.iii) sample of origin. (c) Representative annotated AY and AO sections, coloured by tissue and by Leiden cluster identity. (d) Correspondence between clusters and the 15 tissues, as the fraction of overlapping cells. (e) Colour scales for (d) and (f). (f) Marker gene expression across the annotated tissues; rows are tissues (Metaxylem to Epidermis) and columns are marker genes grouped by tissue. Colour shows relative expression, the mean expression of each gene per tissue min–max scaled from 0 to 1 across the 15 tissues. Asterisks denote significant enrichment of a gene in that tissue, tested by a one-versus-rest Wilcoxon rank-sum test with the area under the receiver-operating curve derived from the U statistic, retaining genes with AUC ≥ 0.70 and a Benjamini– Hochberg-adjusted p < 0.05 (* p < 0.05, ** p < 0.01, *** p < 0.001). Markers were derived by ranking each tissue the genes that best distinguished it from all others and retaining those above the AUC and adjusted-p thresholds. Wheat genome version 2.1 identifier background denotes the source of the marker: blue, identified from the anatomical annotations in this study; light yellow, wheat root atlas (Ke et al., 2025); darker yellow, wheat atlas marker validated across species; light green, atlas marker overlapping an annotation marker from this study; dark green, cross-species atlas marker overlapping an annotation marker, purple boxes show overlap between cell type and expected marker gene.

### Functional enrichment

Over-representation of MapMan bins (Mercator4 v6; Schwacke et al., 2019) and Gene Ontology terms was tested by Fisher’s exact test with Benjamini–Hochberg correction, with fold enrichment calculated relative to the expected proportion. To limit the influence of very lowly expressed genes, enrichment used candidate sets against a background restricted to a mean expression of at least one count per million; terms were called enriched at an adjusted p < 0.05.

### Root hydraulic modelling

Root prototypes were defined by varying the longitudinal distribution of metaxylem number and cross-sectional area along a 70 cm wheat crown root of 28 segments (2.5 cm each). Individual vessel cross-sectional area took one of three values, 1,200, 4,000 or 8,000 µm² (the 12.5th percentile, median and 87.5th percentile of the measured distribution; narrow N, medium M and wide W), and metaxylem file number one of 7, 5, 3 or 1. Prototypes ran through all combinations of initial and final file number and of initial and final area class, at one of three transition rates (one file lost per 2.5, 5 or 10 cm) and beginning either at the root base or after 35 cm, giving 405 prototypes. Root radial conductivity (k_r, m MPa⁻¹ s⁻¹) was modelled with GRANAR–MECHA (Heymans et al., 2020). A reference crown-root anatomy was reconstructed in GRANAR from the median measured parameters (root cross-sectional area 440,000 µm², stele area 64,000 µm², six cortical layers) and coupled with MECHA (Couvreur et al., 2018) using the default subcellular parameters (Table S3). A single longitudinal k_r profile was applied to all prototypes, a high value for the apical 2.5 cm and a low value for the remaining mature root, corresponding to contrasting apoplastic-barrier scenarios (Table S3) and falling within the range reported for wheat (Baca Cabrera et al., 2026; Bramley et al., 2009). Segment axial conductance (k_x, m⁴ MPa⁻¹ s⁻¹) was calculated for each segment from metaxylem number and vessel area by the Hagen–Poiseuille equation (Tyree & Zimmermann, 2002), adjusted for axial hydraulic limitation (Barry et al., 2025). Whole-root conductance was simulated with MARSHAL (Meunier et al., 2020) on a 30-day-old, 70 cm crown root and its laterals generated in CPlantBox (Giraud et al., 2023) and parameterised from the hydroponic measurements (Table S4), with crown-and lateral-root parameters taken from the prototype parameterisation and previous studies respectively. Outputs were whole-root-system conductance (K_rs), segment xylem pressure (MPa) and axial water flow (J_xl).

#### Statistics

Anatomical data (Figures 1 and 2) were analysed in GraphPad Prism v10.4: differences between root classes at each axial position by two-way ANOVA with Tukey’s multiple-comparison correction; the decline in metaxylem file number and complexity along the axis by one-phase exponential decay fits; and relationships between metaxylem area, file number and stele area by Pearson correlation. Candidate gene expression (Figure 3) was compared in GraphPad Prism v10.4 by one-way ANOVA with Tukey’s multiple comparisons across the four sample groups, and by a two-sided Mann–Whitney U test between young and old adventitious cells within each tissue.

## Results

### Metaxylem anatomy converges between adventitious and seminal roots in wheat

Cross sections of young adventitious roots (grown in soil mesocosms) reveals that their vascular anatomy is distinct from comparable positions in seminal roots, particularly by the significant difference in metaxylem file number (Figure 1c.i, c.ii, d) (Table S5). This is apparent both in the older basal sections of roots (Figure 1a.i-iv, b.i-iv) and immediately adjacent to the root tip at comparable distances from the base of the root (Figure 1c.i, c.iii). As cereal roots do not undergo secondary growth, the vascular anatomy observed along the mature root reflects the developmental state of the apical meristem at the time each segment was produced. This enables us to reconstruct developmental changes in metaxylem patterning from mature root anatomy. Sampling at approximately 2.5 cm from the root base, adventitious roots had on average 5 metaxylem files in the stele, whereas seminal roots had only a single centrally positioned file. As adventitious roots grew longer, the number of metaxylem files decreased rapidly, converging on the seminal root phenotype within 30 cm of the root base, while seminal roots maintained their single file throughout development. Metaxylem file number differed significantly between root classes when comparing the segments taken in the first 30 cm, and within the adventitious roots the presence of significant difference in file number in segments and any distal segment ceased between segments after 20 cm. These observations reveal a previously undescribed developmental transition in which anatomically distinct root classes converge towards a common ‘unipolar’ vascular architecture.

This convergence in metaxylem number was not accompanied by a significant change in the total cross-sectional area of metaxylem files or in stele along the root axis in either seminal or adventitious roots. At all positions these values differed significantly between root classes. There was a significant and strong correlation between total metaxylem cross sectional area and stele area in both seminal (r=0.862) and adventitious roots (r=0.956), whereas the number of metaxylem files weakly correlated with stele area (r=0.377). This indicates that while total metaxylem cross sectional area and stele area may be developmentally coupled, the number of files of metaxylem may be a consequence of separate developmental factors and not a process resulting from allometry. Adventitious roots were significantly more complex than seminal roots in the four most basal segments (Figure 1d), and both classes reduced significantly in their mean complexity (normalised perimeter-to-area ratio) along the root axis. Sections within the first 20 cm of adventitious root and the 15cm of seminal root segments showed significant difference when compared to distal sections, then from 25 cm onwards no differences were present between or within root classes. The conjugation of these datasets, sampling at maturity and near the meristem, across different root classes and ages reveal a distinct developmental phase in the root tip in which metaxylem pattern changes with developmental time.

### Root vascular remodelling exists in temperate cereal crops

To determine whether developmental vascular remodelling is unique to wheat or represents a broader feature of cereal root development we quantified metaxylem pattern across many major cereals including oat, rye, barley, several genotypes of wheat across different environments, brome, sorghum, and maize. Remarkably, the same progressive reduction in metaxylem file number along the adventitious root axis was observed in every temperate species. This was reproducible across hydroponic culture, soil mesocosms, and field grown wheat; indicating that vascular remodelling is not an artefact of growth conditions, but reflects an intrinsic and conserved developmental program which leads to convergence between seminal and adventitious root anatomy (Figure 2). This was confirmed by two-way ANOVA, through discovery of significant effects of position along axis, root class, and segment by class interactions, in wheat, barley, rye, oat, and brome.

We extended our studies to also include field grown plants with roots extracted from soil cores as discontinuous sample positions. Consistent with our controlled-environment experiments, no root with more than one metaxylem file was found in samples collected from below 30 cm in any sampled wheat genotypes in our field trials. In contrast, sorghum maintained multiple metaxylem files in all segments across the whole sampled soil profile, reinforcing our conclusion that adventitious root metaxylem pattern remodelling differs fundamentally between the pooideae and panicoid grasses surveyed, regardless of growing conditions.

### A cell-resolved spatial atlas for wheat root tips resolves tissue identities

Having established that vascular remodelling is a conserved developmental programme in temperate cereals, we next sought to identify the molecular changes accompanying this transition. We generated whole-transcriptome spatial transcriptomic profiles (untargeted polyA-captured on the chip’s oligo-dT array) of young and old adventitious root tips using Stereo-seq (Figure 3a), and assigned each encircled capture point (arrayed in a 500 nm grid) to an individual cell of one of fifteen tissue classes with an exclusion zone two capture points thick between cells. This was achieved using the co-registered cell wall image, and provided 6,685 annotated cells, with 4 root tips sections for both the young and old root tips (Figure 3c). Following unguided harmonisation of these cells, this cell pool showed no strong chip or root piece specific separation (Figure 3b.iii). Unsupervised clustering of the 6,685 annotated cells (Leiden resolution 1.4) produced clusters that resolved the major tissue types and closely mirrored the manually assigned identities (Figure 3b.ii).

The clustering highlighted similarities between certain cell types. The clustering strongly validated the assignment of the hypodermis annotation as separate to the rest of the cortex, showing very few hypodermal cells outside cluster 6. The separately annotated cortical cell file layers, (Cortex 1 to 5) predominantly grouped together in cluster 0 (Figure 3d), however there was overlap between the innermost cortical cell file layers (1 and 2) and the endodermis in cluster 4 (Figure 3d). Xylem and metaxylem clustered together in cluster 9, with all metaxylem falling into this cluster, though a small proportion of xylem also appeared in other adjacent clusters. Remaining vascular tissues clustered less cleanly, though both phloem and phloem pole cells peaked in fraction of assigned cells uniquely in their own clusters (7 and 10 respectively). Stele cells neighbouring metaxylem files were assigned their own identity in the manual annotations, however these distributed across clusters 2 and 5 along with the stele parenchyma (Figure 3d).

### Tissue marker genes confirm identity assignment and reveal root tip age differences

Having shown the assigned cell types are transcriptionally distinct, we proceeded to validate these identities using established markers and sought novel markers for our additional cell types. All manually annotated tissue types could be confirmed through peak expression of marker genes from literature in the expected tissue. Several markers from literature also showed high (and in some cases peak) expression in other tissues, affirming the importance of multiple references; however, this challenge was limited to select pericycle and endodermal genes as expected from the cluster overlaps, which had no bearing on downstream analysis. Marker genes for metaxylem were highly specific to the appropriate cells, as were xylem markers, being clearly absent from the other group despite the similarities at clustering. We also confirmed that these marker genes were stable between age groups, showing no significant fold change in the target tissue between young and old root tips.

From our data set we were able to derive candidates for cell type specific markers. Several of these overlapped with marker genes identified in the single cell and spatial wheat root atlas produced by Ke et al. (2025) (Figure 3e, Table S6) (atlas markers in yellow, cross species bright yellow, and green where overlapping with markers identified in this study). Several peroxidases appeared as metaxylem markers in our dataset, which were highly metaxylem specific. In combination, the clustering and marker gene expression analysis strongly support our cell resolved spatial annotations; these also allow identification of redundant annotations or tissue regions containing previously underexplored heterogeneity.

Across the two transcriptomics experiments, expression of 82,074 genes were quantified. From the bulk RNA-seq we identified genes differentially expressed (DESeq2, Benjamini-Hochberg-adjusted p<0.05) between roots contrasting in the number of metaxylem that were being produced at the tip (Table S7). This was done using the contrast groups Adventitious Young versus Adventitious Old (AY/AO), and Adventitious Young versus Seminal Young (AY/SY), with Adventitious Old versus Seminal Old (AO/SO) and Seminal Young versus Seminal Old (SY/SO) as exclusion filters to remove genes that relate primarily to differences in age, phenology, or root class rather than metaxylem file number. Taking just the gene set where the direction of differential expression was concordant in both of the target contrast groups gave 5,093 genes we associated with metaxylem number. This set was then filtered using the spatial data and restricted to genes that were more highly expressed in the pooled metaxylem than in a pseudobulk pool of all other tissues which brought our set down to 1,255 genes of interest. We then filtered this list again, looking for genes that were concordant in direction of change from the many to one bulk pool, and between the young and old spatial pseudobulks (yielding 739 genes). This list was then filtered for genes that showed their peak expression in metaxylem, giving 448 genes of interest. A more stringent filter was also applied at the start, keeping only genes that were also significantly differentially expressed in the spatial pseudo bulk looking for genes that were differentially expressed between chips AY and AO chips. After filtering for concordance with the bulk RNA-seq and applying the other described filters this yielded five genes of interest, one upregulated in young adventitious root tips relative to all other sample groups (Figure 4d), and four downregulated, one given as an example in Figure 4f. Functional enrichment varied with filtering stage and root tip age (Figure 4b), with transcriptional co-regulation enriched in the young metaxylem-specific set and cell-cycle organisation and microtubular network terms in both old root tip sets.

**Figure 4.**
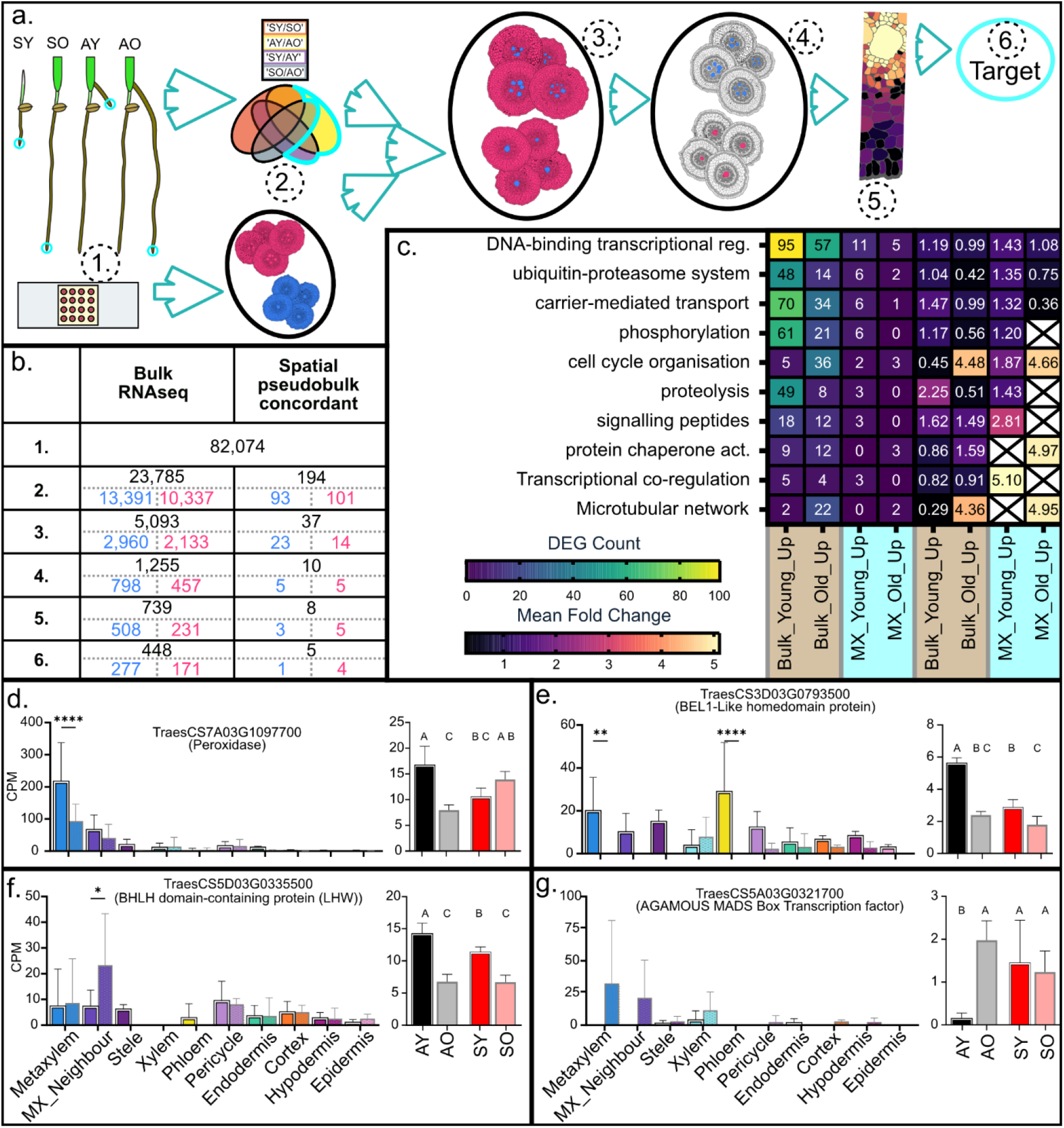
Integrated filtering of the bulk and spatial transcriptomes nominates candidate regulators of metaxylem convergence. (a) Filtering pipeline: root tips of the four groups (seminal young, SY; seminal old, SO; adventitious young, AY; adventitious old, AO) were profiled by bulk RNA-seq and Stereo-seq and filtered through six steps (1 to 6, corresponding to the rows of b). (b) Genes retained at each step, for the primary bulk-anchored set and the parallel spatial-pseudobulk-concordant set; up-and down-regulated in blue and pink. Step 1, genes detected in both platforms (82,074); 2, differentially expressed in the bulk (DESeq2, Benjamini-Hochberg-adjusted p < 0.05); 3, concordant across the two target contrasts (AY/AO, AY/SY) excluding SY/SO, AO/SO; 4, more highly expressed in pooled metaxylem than in a pseudobulk of all other tissues; 5, concordant between the bulk and the spatial metaxylem pseudobulks; 6, per-cell expression peaks in metaxylem. (c) MapMan enrichment of the bulk and metaxylem (MX) young-up and old-up sets; left, gene count per bin; right, mean fold change; crosses, bins with no genes in the set. (d to g) Four candidates, per-tissue CPM in the spatial data (AY left with white border, AO right) and per-group CPM from the bulk; (d) a class III peroxidase (TraesCS7A03G1097700), (e) a BEL1-like homeodomain factor (TraesCS3D03G0793500), (f) an LHW-like bHLH factor (TraesCS5D03G0335500) and (g) an AGAMOUS-type MADS-box factor (TraesCS5A03G0321700). Bars, mean ± SD (n = 4). Significance: (d–g, left) two-sided Mann-Whitney U test comparing young and old adventitious cells within each tissue; (d–g, right) one-way ANOVA with Tukey’s multiple comparisons, shared letters denoting groups that do not differ. * p < 0.05, ** p < 0.01, **** p < 0.0001

The most strongly metaxylem specific genes in the spatially cell filtered set of 488 genes were peroxidases (Class III, which catalyse lignification of vessel walls), comprising the whole top six (Figure 4d), several of which were identified as marker genes in the earlier analysis (Figure 3e). Though their overall and per cell expression changed with root tip age, they were more highly expressed in metaxylem than any other cell type in both conditions. Several bHLH transcription factors appeared in the 448 set list, including an orthologue of Lonesome-Highway (LHW), although closer inspection revealed age effects in the seminal roots and a less clear spatial expression pattern, it is notable this candidate with a well characterised role in determining xylem pole number in *Arabidopsis* and rice was surfaced. In the five gene subset with significant changes in expression in the spatial pseudobulk were two genes with functional annotations relating them to meristem maintenance (Figure 4e,f). These were a BEL1-Like homeodomain protein (a positive regulator of meristematic cell division), and an Agamous MADS-box Transcription factor, a repressor of meristematic cell division, more highly expressed in the poly-metaxylem file tip state and mono-metaxylem file tip state respectively. Genes homologous to or bearing functional annotations of Forked or Forked-like family members appeared in the unfiltered bulk list, but had a similar expression pattern in seminal as well as adventitious roots and were expressed more strongly in xylem and stele parenchyma than metaxylem.

This conservative filtering strategy prioritizes specificity over breadth, potentially excluding regulators with broader spatial expression or those acting upstream of root class and meristem age. Despite these stringent criteria, the remaining candidate dataset captured significant transcriptional shifts across successive stages of metaxylem determinacy, including meristematic maintenance, cell division, cell elongation, and cell wall modification. Rather than pointing to a single master regulator, these results demonstrate a multi-layered regulatory program governing the metaxylem transition, even within the narrow developmental window captured by the spatial data. Consequently, while these cell-type filters may not capture the entire regulatory network, they successfully isolate the core parallel processes driving this anatomical shift, providing a robust foundation for future functional validation.

### Metaxylem number and vessel size shape whole-root conductance and xylem pressure

To assess the functional consequences of metaxylem file remodelling, we simulated the hydraulic properties of roots with different patterns of metaxylem change under different basal and distal anatomies, using GRANAR (Heymans et al., 2020) and MECHA (Couvreur et al., 2018) to estimate realistic radial uptake values and MARSHAL (Meunier et al., 2019) to integrate positional values. Using this integrated modelling framework allowed varying of only metaxylem patterns along the axis and no other traits in order to target the specific functional impacts of metaxylem file patterning exclusively. Root prototypes were digitally constructed, contrasting in basal and distal metaxylem file number, initial and final mean metaxylem cross-sectional area, transition rate (distance in cm for a mx file to be lost) and root length. Here we focus on the contrast between anatomically uniform reference prototypes and several root morphologies whose longitudinal transition patterns match the observed phenotypes. At the level of a single cross section, axial conductance (Kx) is proportional to the sum of the fourth power of the vessel radii, and can therefore be dominated by the width of the individual files in the segment, so a change to fewer, larger vessels has a predictably multiplying effect on segment conductance. The behaviour of the root as a whole is less intuitive: the axial conductance of each segment governs all the water that must pass through it toward the shoot, so a change in the basal region, through which the entire root hydraulic movement travels, weighs far more heavily than the same change near the tip, and capturing this requires integrating conductance along the whole root. This is also due to the additive effects of axial limitations, meaning for a specific axial conductance, additional root length will not affect whole root conductance (Meunier et al. 2017, Baca Cabrera et al. 2024).

This integration showed that the whole-root cost of a progressive loss of metaxylem files depends on how that loss occurs. Relative to the matched unchanging prototype, scenarios in which the metaxylem file pattern transitioned to a wider single central file, in the manner seen in wheat adventitious roots, typically increased whole-root conductance (Krs). This increase occurs even when seven narrow files converging to a single medium area file (7N:1M), reduces the final metaxylem area to less than half of the starting area. In a further scenario, where five medium vessels with a total area of 20,000 µm² transitioned to a single vessel of 8,000 µm², whole-root conductance was maintained at 98% of the unchanging prototype, despite the axial conductance of the post-transition segment being only 80% of that of the starting condition. Conversely, in the seminal root prototypes, which start and end with a single vessel and change only in its cross-sectional area, reducing that area by 70% or increasing it by 100% relative to the 4,000 µm² unchanging scenario, results in an eleven-fold decrease and a four-fold increase in Kx respectively, but only a 65% decrease and a 31% increase in Krs (Figure 5, Table S8).

**Figure 5.**
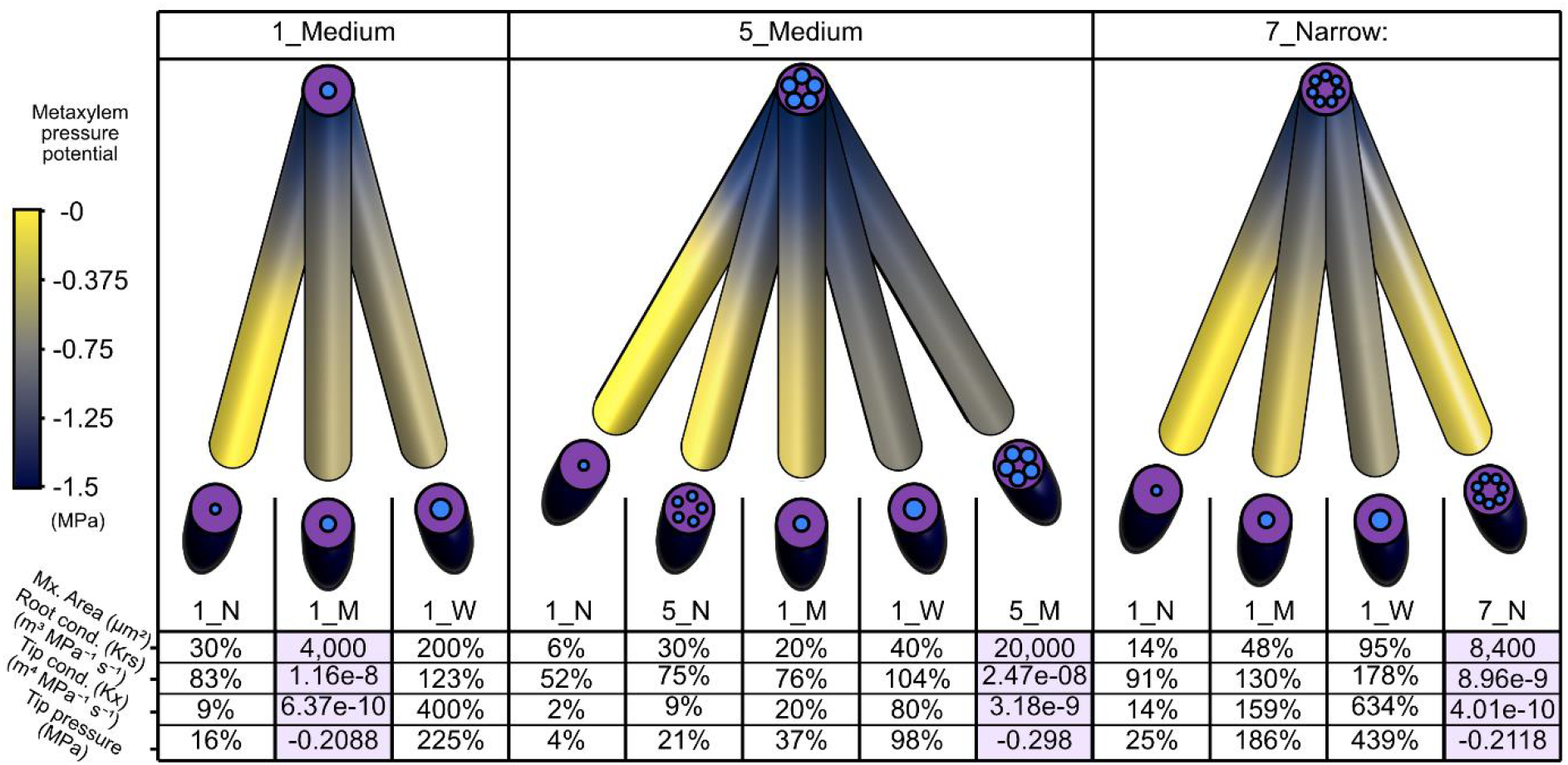
**Metaxylem file number and vessel size shape whole-root conductance and xylem pressure**. Hydraulic properties of modelled wheat root prototypes of defined metaxylem configuration, simulated with GRANAR, MECHA and MARSHAL. Each configuration is named by metaxylem file number and cross-sectional area class (N, narrow; M, medium; W, wide); Prototypes share one of three basal anatomies (1_Medium, 5_Medium and 7_Narrow) against which the others are compared. Each prototype radiates from the shoot junction and is coloured along its length by the metaxylem hydraulic pressure potential (MPa; colour bar), with tip stele anatomy metaxylem shown in cross section at each end. The table below each panel gives the total metaxylem cross-sectional area, whole-root conductance (Krs), tip conductance (Kx) and the metaxylem pressure potential at the tip; each is expressed as a percentage of the reference anatomically unchanging configuration, with the reference itself (highlighted) given as absolute values

These divergent hydraulic responses demonstrate that whole-root conductance can either buffer or magnify the localized impacts of vascular remodelling depending on the specific geometry of the transition. This indicates that anatomical continuity along the longitudinal axis is an independent and highly impactful property determining root hydraulic function, decoupled from simple cross-sectional area metrics.

## Discussion

### A heteroblastic phase change in root apical meristems of temperate cereals

The metaxylem anatomy of cereal adventitious roots is not fixed at root genesis, but in the developing region is progressively remodelled during root growth until a final production state of a single central file is reached. As this pattern is largely constitutive in seminal roots, vascular patterning is convergent in the axial root classes. Viewed along the root axis in a progressively distal view, the metaxylem undergo a confluence, though when viewed in the direction of xylem sap flow and from the apical meristem towards older tissue this reads as a furcation (Alemán-Sancheschúlz et al., 2026). At maturity, it is the case that the basal, older root segments typically contain more files of metaxylem than distal segments, but as we show through our time series studies, this is better interpreted as the distinct vascular anatomy as produced by a ‘young’ root tip rather than a consequence of tissue age. For these reasons we consider the metaxylem pattern along mature cereal adventitious roots a feature that is more readily interpreted in distance from the point of organogenesis at the root junction as opposed to relative to the root tip.

Given how fundamental metaxylem function it is striking that this phenomenon is not more widely documented, however this speaks to the technical challenges of recovering intact lengths of adventitious roots. Several studies have however presented data where this transition is visible (Wu et al., 2011; Ouyang et al., 2020; Baca Cabrera et al., 2026); mostly focusing on the changes in lignification and secondary barrier formation along the root axis. Insights into species not included in our surveys can be gleaned from these studies, and interestingly present examples of contrast to the pattern we observe in the cool season grasses. Rate of vascular change as a trait appears to vary within wheat, and shows variation within a panel of winter wheat cultivars released over 120 years of breeding (Baca Cabrera et al., 2026; Behrend et al., 2025). This difference between cultivars was heritable and showed a gradient in appearance that aligned with cultivar release year, with more modern cultivars showing a greater extent of vascular remodelling within the same length of root sampled.

What defines this as an anatomical developmental transition rather than a gradual loss of meristematic proliferative potency is that after this transition occurred, in no cases did the root cease to grow at the end of this phase of change, or did there cease to be a central metaxylem file in the root, as can be seen in lateral roots (Shishkova et al., 2008). In all of these species in which the distinct remodelling occurred, regardless of the starting condition, or the length of root produced before this transition was complete, the endpoint was the same and irrevocable, and had no clear bearing on final root length.

### Meristem progression drives the meta-xylem-morphoses

The reduction in metaxylem file number reflects a change in what the root apical meristem produces over developmental time, rather than any alteration to a segment once it has matured. Consistent with this, when we imaged root tips across a range of root lengths, the youngest, shortest tips contained the most metaxylem files close to the tip, and this number fell as roots lengthened and tips aged. We can therefore divide adventitious root development into two phases: an initial transition phase following emergence, in which multiple files are present and the vasculature is actively remodelled, and a subsequent stable phase, in which a single central file is produced and maintained.

How the number of metaxylem files produced at the meristem, and the change in that number, is regulated remains a key question that will require further exploration. A next step will be to clarify the cellular mechanism of this remodelling: whether multiple metaxylem initials relocate to become neighbouring fuse, and or are eliminated, whether a single initial divides and branches (Miki et al., 2026; Saito et al., 2019) before differentiation in the early phase and ceases to branch later, or whether a combination of these occurs, with multiple initials both changing their division pattern and being eliminated. Given the large spatial distance over which this transition occurs relative to the size of vascular cells pre-elongation, resolving this will require high-resolution three-dimensional anatomical surveys, or high frequency time-point sampling with lineage tracing of cell files in the root tip. What is clear is that this process must be tightly regulated to preserve hydraulic continuity and root function (Sawchuk et al., 2013; Etchells et al., 2016; Turco et al., 2019). We suggest that this behaves as a developmental phase transition of the root meristem, comparable to the phase transitions long recognised in shoot meristems, such as the switch from vegetative to reproductive (Huĳser and Schmid, 2011) growth, but expressed through vascular anatomy rather than organ identity. Such transitions are well characterised in shoots, yet remain little explored in roots and even less so in grasses (Brady et al., 2006, 2007; Yang et al., 2024; Rodriguez-Alonso et al., 2018). Developmental regulators can similarly be conserved or recruited to control distinct developmental outcomes across grasses (Klein et al., 2022; Xiao et al., 2022; Gallagher et al., 2023).

To target these changes, we profiled the transcriptomes of root meristems producing contrasting vascular anatomies over the course of development, using bulk RNA-seq to define a candidate pool and spatial transcriptomics to filter it to genes with focused metaxylem expression and concordant change. This conservative, two-technique strategy isolates a smaller set of higher-confidence targets than either method alone, to prioritise for functional study in transformed or mutagenised populations. We devised a strategy that draws on the complementary strengths of the different transcriptomic platforms. Spatial transcriptomics provides excellent spatial resolution and allows direct annotation of cell types. It is, however, resource-intensive and low in capture rate (De Veylder et al. 2026), and because metaxylem cells are scarce, especially in the mono-file state, our spatial data alone struggles to provide the replication needed to serve as our main discovery platform. Bulk RNA-seq is complementary, robust and well-replicated but not cell-type specific, and so carries the differential-expression discoveries, while the spatial data validates and filters the candidates. This combines the stringency of the spatial with the robustness of the bulk, an approach that should generalise to other low-abundance cell types. Filtering against the spatial data often increased the mean fold change of the enriched functional groups (Figure 3B), indicating that it not only shortens the candidate list but enriches for the more strongly varying genes of interest.

The candidates fall into two groups, the first relating directly to the vessels. An LHW-like bHLH transcription factor was present among the metaxylem-associated genes, of interest given the established role of LHW in setting xylem pole number in *Arabidopsis* (De Rybel et al., 2013; Ohashi-Ito et al., 2024), and its disruption of metaxylem file reduction in rice (Reeger et al., 2021). Several peroxidases were among the most strongly metaxylem-specific genes, most plausibly reporting metaxylem differentiation rather than driving the change in file number due to their role in catalysing the oxidative polymerisation of mono-lignols during cell wall lignification (Marjamaa et al., 2009). Their per-cell expression also fell from young to old tips, which is itself informative: the signal is strongest when metaxylem files are most numerous, so bulk data alone might have ascribed it simply to a greater number of metaxylem cells, whereas the cell-resolved data show a genuine change in expression per cell. The second group relates more to the meristem. Among the genes differentially expressed between young and old adventitious tips were a BEL1-like homeodomain factor and an AGAMOUS-type MADS-box factor, both with a predicted role in meristem maintenance and determinacy (Kumar et al., 2007; Cao et al., 2024).

AGAMOUS is known for specifying a shift in identity at the floral meristem (Bartlett, 2017; Abraham-Juárez et al., 2020; Tapia-López et al., 2008; Montiel et al., 2020), and its increased expression in older tips is consistent with a meristem progressing towards a more determinate, single-file output. Together these analyses do not immediately resolve the regulatory network underlying the transition (Brady et al., 2011), but they isolate candidate processes and, alongside the validated cell-resolved atlas of the wheat root tip, establish wheat as a tractable system in which to dissect developmental phase transitions in roots.

### Evolutionary divergence of the transition across grasses

The transition is not universal among grasses. It occurred in every genotype of the temperate species we surveyed, wheat, barley, rye, oat and *Brachypodium*, but was absent in the PACMAD cereals maize and sorghum, the adventitious roots of which retained multiple metaxylem files along their entire length and remained anatomically distinct from their seminal roots. Basal metaxylem file number and area can differ with species, genotype and stem node of origin in both temperate and tropical cereals, and have been the focus of many studies linking root anatomy to plant physiology and agronomic (Strock et al., 2021; Klein et al., 2024; Affortit et al., 2024) performance. Basal anatomy has been shown to be an important trait, but our results indicate that in many species sampling at this position is insufficient.

Why the transition is present in the temperate grasses surveyed but not the tropical ones is intriguing, and plausibly relates to adaptation to their respective ecological niches; the contrast may in part reflect the different water-use and conservation strategies associated with C3 and C4 (Gowik and Westhoff, 2011) photosynthesis. The BOP clade offers further contrasts: though changes in metaxylem number do occur, neither rice nor intermediate wheatgrass show the rate or extent of remodelling (Ouyang et al., 2020; Clément et al., 2022) seen in the other members we surveyed, which may reflect the distinct demands of a semi-aquatic habit and a perennial lifecycle respectively. The trait may therefore segregate along several axes, including clade, photosynthetic type, lifecycle and adaptation to water availability (Kadam et al., 2015) or other edaphic gradients such as soil density, nutrient availability and temperature (Zobel, 1992; Peralta Ogorek et al., 2025). Alongside a fuller characterisation of the developmental landscape of this remodelling, surveying more diverse grasses adapted to contrasting environments should help resolve its regulation and function.

### Functional consequences

The remodelling also carries consequences for the hydraulic architecture of the root. Modelling of the measured anatomy indicated that converging on a single, enlarged central vessel is an efficient way to maintain whole-root axial conductance (Hacke et al., 2001; Tyree and Zimmermann, 2002) as the root remodels, because the outcome depends more on the configuration through which metaxylem area changes than on the amount of area lost; concentrating area into fewer, larger vessels preserves or increases conductance, whereas an equivalent loss of area spread across an unchanged number of files does not. Because axial conductance integrates along the whole root, holding one configuration along the entire axis would compound its costs (Meunier et al., 2017; Baca Cabrera et al., 2024), whereas the graded transition matches the supply capacity of each region to the demand placed on it: multiple files where the cumulative flow from the whole root passes basally, converging to a single efficient vessel distally where flow is lower.

Having two distinct vascular states may balance several competing demands over the life of the root, in particular a trade-off between robustness and efficiency. A single wide conduit is more vulnerable to failure through embolism (Harrison Day et al., 2023) than several narrower ones in which the failure of any one removes less of the total conductance; retaining files basally, where the consequences of failure are greatest, while converging distally as tension rises places robustness and efficiency where each is most useful. The risk of failure at greater depths may also be reduced, being upstream of the additive longitudinal pressure gradient produced by friction (Bouda et al., 2018) of water transport. Thus, rather than responding plastically to the local soil hydraulic environment, developmental remodelling of metaxylem patterning may preconfigure different regions of the root for the hydraulic conditions they are likely to experience later in the plant life cycle. Older basal tissues retained in the upper soil profile maintain multiple conduits, whereas younger tissues extending into deeper soil are produced with a hydraulically distinct vascular configuration. The reduction in vessel area may also (Richards and Passioura, 1989) conserve water, favouring an architecturally distributed root network over a high-conductance one built from fewer roots, and could suit growth through vertically heterogeneous soil, where water, mechanical impedance, temperature and nutrients all change with depth, as too does the intensity of environmental fluctuations (Baumberger et al., 2024; Philipp et al., 2025). There may additionally be a construction cost: for a given area, many small vessels present far more wall surface than one large vessel, since perimeter scales less steeply than area, so consolidating metaxylem may reduce the lignified wall built per unit of conductive area and be more economical in carbon. We treat these ecological and economic interpretations as provisional, since our functional evidence is confined to the modelling.

This work uncovers a previously undescribed developmental transition in cereal roots and provides a validated cell-resolved atlas of different states of the adventitious root tip. This incident of change in vascular patterning is developmentally novel and, on the evidence of the modelling, physiologically consequential, both in understanding the consequence of vascular change in adventitious roots, and in understanding the functional differences of adventitious and seminal roots. Consideration of this phenomenon will be essential in future of how vascular pattern is built and maintained across the life of cereal roots and the consequences thereof.

## Author Contributions

D.H.J.: conceptualization, data curation, formal analysis, investigation, methodology, visualization, writing-original draft; M.Sö.: data curation, formal analysis, investigation, methodology, visualization, writing-review & editing; D.D.: investigation, resources, methodology; C.T.: investigation, resources, methodology; J.B.C.: investigation (hydraulic modelling), formal analysis, data curation, methodology, writing-review & editing; A.W.: conceptualization, data curation, methodology, writing-review & editing; M.M.: investigation; M.Sc.: investigation, resources, methodology, writing-review & editing; D.Z.: investigation, writing-review & editing; R.G.: investigation, writing-review & editing; G.L.: writing-review & editing; B.P.: writing-review & editing; V.C.: writing-review & editing; M.B.: writing-review & editing; D.R.: project administration, funding acquisition, writing-review & editing; H.M.S.: conceptualization, funding acquisition, supervision, project administration, methodology, writing-review & editing. All authors reviewed and approved the final manuscript.

## Supplementary Material

Table S1 List of genotypes used in this study

Table S2 Mean, standard deviation and n for each species, root class and axial position, for all measured vascular traits (Figures 1 and 2).

Table S3 Subcellular hydraulic parameters and apoplastic barrier configuration used for MECHA simulations.

Table S4 Root architectural and hydraulic parameters used for whole-root hydraulic simulations.

Table S5 Whole-root-system architecture of hydroponically grown wheat, by node and axial position (mean, SD, n).

Table S6 Cell-type marker genes used to validate the spatial annotation, with the tissue each marks, its source and discrimination statistics (Figure 4).

Table S7 Bulk RNA-seq: raw read counts per sample, DESeq2 contrasts, and the candidate-filtering stage each gene reaches (Figure 3).

Table S8 Hydraulic simulation outputs for all 405 root prototypes (Figure 5).

## Funding

This study was supported by the Grains Research and Development Corporation (GRDC) through grant “Root structure and function traits: Overcoming the root phenotyping bottleneck in cereals (UOQ2312-009RTX) (D.H.J), the Leibniz Association (Resilient Roots to H.M.S.), and the European Union (ERC, 101162856, FATE) (H.M.S.). JCBC is funded by Deutsche Forschungsgemeinschaft (DFG, German Research Foundation) (SFB 1502/1–2022, Projektnummer: 450058266. DZ is the recipient of an Australian Research Council Early Career Industry Fellowship (IE230100179, “Drought tolerance in sorghum: the roots of the solution”) and funded by Australian Government. VC is a Research Associate of the Belgian Fonds de la Recherche Scientifique (FNRS), co-funded by the European Union (ERC grant 101043083). The views and opinions expressed are, however, those of the author(s) only and do not necessarily reflect those of the European Union or the European Research Council. Neither the European Union nor the granting authority can be held responsible.

## Conflicts of interest

The authors declare no conflicts of interest.

## Data availability statement

Data available in article supplementary material and upon request.

## Supporting information

Supplemental_hydroponic_parameters

Supplemental_Datasheet

## Notes

### Competing Interest Statement

The authors have declared no competing interest.

