## Supplemental_hydroponic_parameters for "Meta(xylem)morphoses: Root vascular patterning is dynamic and convergent in temperate grasses"

**Table S1**: Subcellular hydraulic parameters and apoplastic barrier configuration used for MECHA simulations. Default subcellular hydraulic parameter values from Couvreur et al. (2018, Table 2) were used. The apoplastic barrier configuration was defined according to reported longitudinal anatomical gradients in wheat roots (Bramley et al., 2007, 2009; Baca Cabrera et al., 2026).

| **Parameter** | **Value** | **Unit** |
| --- | --- | --- |
| Cell wall hydraulic conductivity | 2.8 ×10^-9^ | m^2^ MPa^-1^ s^-1^ |
| Hydraulic conductivity of suberized and lignified cell walls | Hydrophobic | - |
| Plasma membrane intrinsic hydraulic conductivity (*k*_m_) | 3.5 ×10^-8^ | m MPa^-1^ s^-1^ |
| Contribution of aquaporins to plasma membrane hydraulic conductivity (*k*_AQP_) | 5.0 ×10^-7^ | m MPa^-1^ s^-1^ |
| Conductance of plasmodesmata per unit membrane surface (*k*_PD_) | 2.4 ×10^-7^ | m MPa^-1^ s^-1^ |
| Protoplast permeability (*L*_pc_) | 7.74 ×10^-7^ | m MPa^-1^ s^-1^ |
| Apoplastic barrier (tip) | Endodermis with Casparian strip | - |
| Apoplastic barrier (mature root) | Fully suberized endodermis | - |

**Table S2**: Root architectural and hydraulic parameters used for whole-root hydraulic simulations. Parameters were used to construct the representative wheat root architecture in CPlantBox and perform whole-root hydraulic simulations with MARSHAL. Root hydraulic parameters (*k*_r_ and *k*_x_) were determined from GRANAR–MECHA simulations, metaxylem anatomical measurements, and previous studies (Baca Cabrera et al., 2026; Bramley et al., 2009).

| **Parameter** | **Value** | | **Determination** |
| --- | --- | --- | --- |
| Root system age (days) | 30 | Hydroponic experiments | |
| Crown root radius (cm) | 0.0375 | Hydroponic experiments | |
| Crown root growth rate (cm day^-1^) | 2.4 | Hydroponic experiments | |
| Crown root apical zone (cm) | 4.8 | Hydroponic experiments | |
| Crown root basal zone (cm) | 2.4 | Hydroponic experiments | |
| Inter-lateral distance (cm) | 0.2 | Hydroponic experiments | |
| Lateral root radius (cm) | 0.014 | Hydroponic experiments | |
| Lateral root growth rate (cm day^-1^) | 0.8 | Hydroponic experiments | |
| *k*_r_ crown root mature tissues (m MPa^-1^ s^-1^) | 5.4 ×10^−8^ | GRANAR–MECHA and previous studies | |
| *k*_r_ crown root tip (m MPa^-1^ s^-1^) | 5.1 ×10^−7^ | GRANAR–MECHA and previous studies | |
| *k*_x_ crown root (m^4^ MPa^-1^ s^-1^) | 5.7 ×10^−12^ – 1.8 ×10^−9^ (segment-specific) | Calculated from metaxylem number and diameter | |
| *k*_r_ lateral root (m MPa^-1^ s^-1^) | 1.3 ×10^−7^ | GRANAR–MECHA and previous studies | |
| *k*_x_ lateral root (m^4^ MPa^-1^ s^-1^) | 1.0 ×10^−13^ | Literature values | |
